# Identification of a Novel Ferroptosis Inducer with Dual Modulatory Effects on GPX4 Activity and Stability

**DOI:** 10.1101/2023.12.22.572948

**Authors:** Jun Wang, Long Liao, Bo Yang, Beiping Miao, Botai Li, Xuhui Ma, Annika Fitz, Shanshan Wu, Jia He, Qianqian Zhang, Shuyi Ji, Guangzhi Jin, Jianming Zhang, René Bernards, Wenxin Qin, Chong Sun, Cun Wang

## Abstract

Ferroptosis is a unique form of intracellular iron-dependent cell death that differs from apoptosis, necrosis, and autophagy. GPX4, an antioxidant defense enzyme, plays a pivotal role as regulator of ferroptosis. Extensive researches suggest that targeting GPX4 holds promise for cancer therapy. However, the current GPX4 inhibitors face challenges due to unfavorable drug-like properties, which hinder their progress in clinical development. In this study, we identified a novel inhibitor called MI-2, demonstrating potent ferroptosis-inducing capacity. Mechanistically, MI-2 effectively inhibits the activity of GPX4 by direct interaction. Furthermore, MI-2 promotes the degradation of GPX4 through its well-established target, MALT1. In multiple cancer models, MI-2 has demonstrated synergistic effects when combined with sorafenib or regorafenib, resulting in enhanced ferroptosis induction. These findings highlight the dual modulatory effects of MI-2 on GPX4 activity and stability, offering a promising starting point for the development of drug-like GPX4 inhibitors with translational potential.

## Introduction

Ferroptosis is a newly identified form of programmed cell death that is characterized by the accumulation of iron-dependent lipid peroxidation and subsequent membrane damage^1^. Unlike other forms of cell death, such as apoptosis or necrosis, ferroptosis relies on intracellular iron levels and lipid metabolism^1,2^. Recent years have witnessed growing interest in studying the role of ferroptosis in tumorigenesis and cancer therapy^3,4^. Certain types of cancer cells, characterized by specific metabolic features, genetic mutations, or imbalanced ferroptosis defense mechanisms, display heightened susceptibility to ferroptosis^5,6^. These findings have sparked interest in exploiting ferroptosis as a potential therapeutic strategy for cancer treatment. Moreover, researchers are exploring combination therapies involving ferroptosis inducers alongside with conventional cancer therapies to enhance the effectiveness of the treatments across various malignancies.

GPX4 plays a crucial role in regulating ferroptosis by reducing lipid hydroperoxides and safeguarding cells against oxidative damage^3^. However, developing effective inhibitors for GPX4 has proven challenging because it lacks a drug-like binding pocket and relies on a nucleophilic selenocysteine residue for its enzymatic activity^7^. Covalent inhibitors, like RSL3, ML162, ML210, and their derivatives (such as JKE-1674) have been developed to target GPX4^3,7,8^. However, these inhibitors face challenges due to poor pharmacokinetic properties and limited selectivity, which restrict their potential for clinical translation^8^. Therefore, it is crucial to develop novel GPX4-targeting agents and gain a comprehensive understanding of the underlying mechanisms of these newly identified inhibitors. This will facilitate the development of more promising therapeutic options based on exploiting ferroptosis for the treatment of cancer.

Liver cancer is a major cause of cancer-related deaths worldwide, but treating advanced-stage liver cancer is challenging due to limited therapeutic options^9–11^. Receptor tyrosine kinase (RTK) inhibitors, like sorafenib and lenvatinib, offer modest overall survival benefits^12,13^. Immunotherapy based options, specifically the combination of atezolizumab (anti-PD-L1 monoclonal antibody) and bevacizumab (anti-vascular endothelial growth factor monoclonal antibody), has shown promise in some advanced liver cancer cases^14–16^. However, many patients with advanced liver cancer still suffer from therapeutic resistance and disease progression. Researchers propose triggering ferroptosis as a strategy to combat established liver cancer in preclinical stages. For example, the combination of donafenib and GSK-J4 has demonstrated synergistic induction of ferroptosis by upregulating HMOX1 expression^17^. Concurrent induction of ferroptosis and blockade of MDSC sensitize liver tumors to immune checkpoint blockade^18^. Additionally, the LIFR-NF-κB-LCN2 axis, which can be targeted, controls susceptibility of liver cancer cells to ferroptosis^19^. Notably, sorafenib, the first-line drug for advanced liver cancer, has exhibited ferroptosis-inducing properties through targeting the activity of SLC7A11, despite some controversy surrounding this conclusion^20,21^. Considering these findings, investigating the potential of harnessing ferroptosis for liver cancer treatment is warranted.

In our study, we identified a small-molecule compound called MI-2 that has the ability to induce ferroptosis in cancer cells. This effect is achieved through the compound’s dual modulatory effects on both GPX4 activity and stability. We not only investigated the underlying mechanisms of these dual modulatory effects but also assessed the synergistic effects of MI-2 in combination with sorafenib or regorafenib across various types of malignancies.

## Results

### Identification of MI-2 as a ferroptosis inducer in liver cancer cells

To assess the potential of leveraging ferroptosis in liver cancer as a therapeutic approach, we initially examined the expression levels of GPX4 in HCC tissues. We observed an upregulation of GPX4 expression in tumor tissues compared to non-tumor tissues at the mRNA level (Extended Data Fig. 1a), and this was corroborated at the protein level through immunohistochemical analysis (Extended Data Fig. 1b). Additionally, within a cohort of 365 HCC patients, those with the highest levels of GPX4 mRNA in their tumors experienced the poorest survival outcomes (Extended Data Fig. 1c). Both pharmacological and genetic inhibition of GPX4 noticeably suppressed the proliferation of liver cancer cells (Extended Data Fig. 1d-f).

Current GPX4 inhibitors, such as RSL3 and ML210, face limitations for *in vivo* applications due to poor bioavailability and low selectivity^8^. In this study, we conducted a compound screening involving 2103 compounds on two ferroptosis-sensitive liver cancer cell lines, with a focus on identifying potential ferroptosis inducers evidenced by the rescue effects of ferrostatin-1 on cell viability (Fig. 1a, b). Our results identified MI-2, EX 527 (selisistat), and AI-10-49 as potential candidate drugs capable of inducing ferroptosis in both cell lines (Fig. 1c, d).

**Figure 1.**
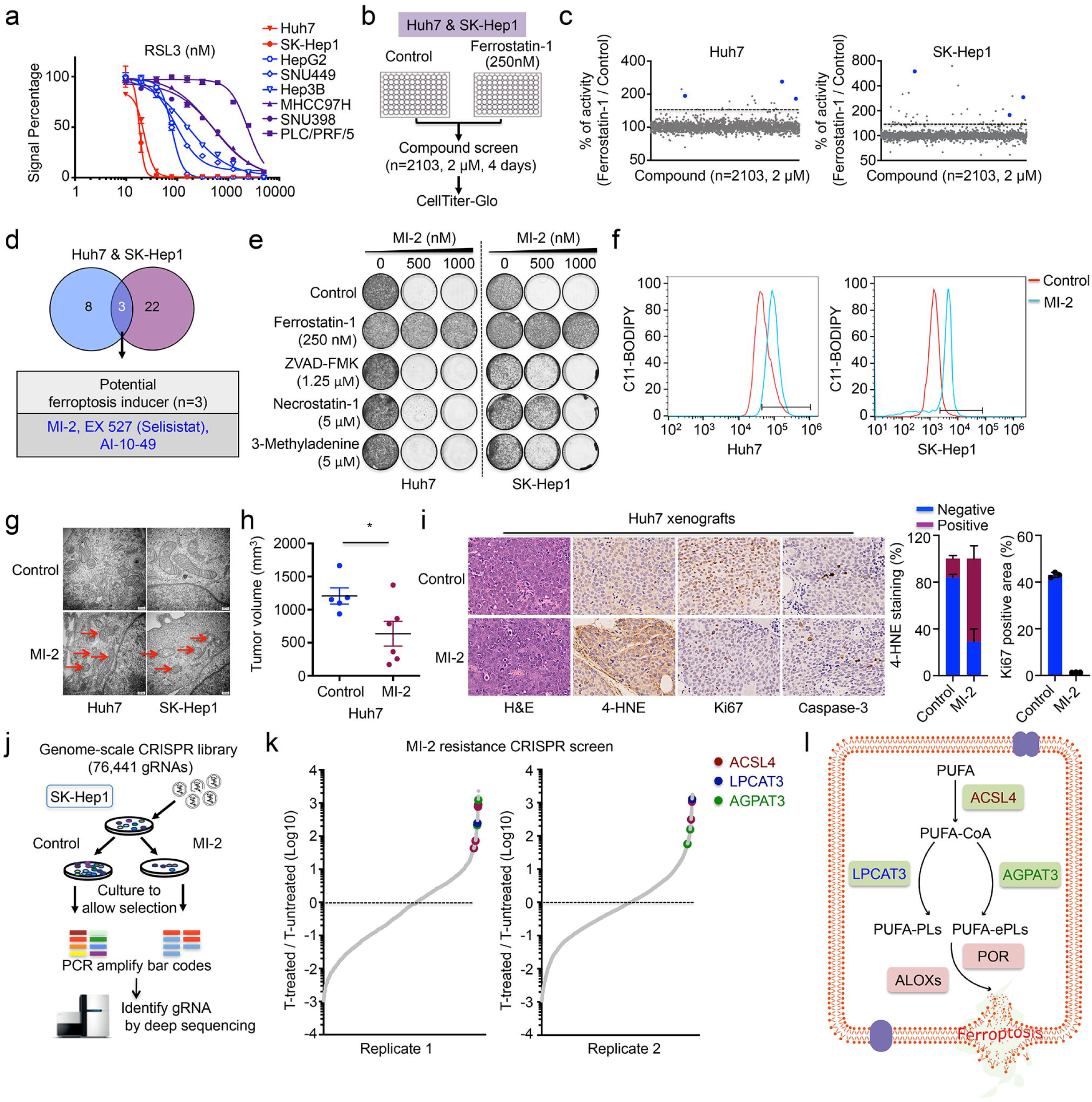
Identification of MI-2 as a ferroptosis inducer in liver cancer cells. **a,** Viability of liver cancer cells lines treated with increasing concentrations of RSL3 for 96 hours. **b,** Schematic outline of the compound screening. Huh7 and SK-Hep1 cells were treated with a compound library (n=2103, 2 μM) in the presence or absence of the ferroptosis inhibitor ferrostatin-1 (250 nM) for 4 days in two replicates. Cell viability was assessed by CellTiter-Glo. **c-d,** A total of three compounds (MI-2, EX 527, AI-10-49) were identified as potential ferroptosis inducers in both Huh7 and SK-Hep1 cells, evidenced by the rescued effects of ferrostatin-1 on cell viability (at least 1.5-fold increase in the group treated with combination of the compound and ferrostatin-1 compared to the group treated with the compound alone). **e,** Colony formation assays were performed on indicated cell lines treated with MI-2 combined with ferrostatin-1, ZVAD-FMK, necrostatin-1, or 3-Methyladenine, respectively for 10–14 days. Ferrostatin-1, the ferroptosis inhibitor; ZVAD-FMK, the apoptosis inhibitor; necrostatin-1, the necroptosis inhibitor; 3-Methyladenine, the autophagy inhibitor. **f,** Huh7 and SK-Hep1 cells were treated with MI-2 (2 μM) and C11-BODIPY (2 μM) oxidation was assessed by flow cytometry. **g,** Ultrastructural analysis of MI-2-treated cells (Huh7 and SK-Hep1) using transmission electron microscopy. Red arrows indicate damaged mitochondria. **h,** Tumor volumes of Huh7 tumor xenografts in BALB/c nude mice were measured following vehicle or MI-2 (20Lmg/kg) treatment for 12 days. \**P* < 0.05. **i,** Representative images of H&E, 4-HNE, Ki-67, and cleaved caspase-3 staining in Huh7 xenografts of mice treated with either vehicle or MI-2. Quantification of 4-HNE and Ki-67 staining from each group (right panels). **j,** Schematic representation of the genome-wide CRISPR-Cas9 screening performed in SK-Hep1 cells. SK-Hep1 cells were infected with a lentiviral genome gRNA library and cultured for 45 days in the absence (untreated group) or presence of MI-2 (700 nM, treated group) in two replicates. gRNA barcodes of untreated or treated samples were recovered by PCR and analyzed by next generation sequencing. **k,** Representation of the relative abundance of the gRNA barcode sequences from the genome-wide resistance screens. ACSL4, LPCAT3 and AGPAT3 were identified as the candidates whose knockouts conferred resistance to MI-2 treatment. **l,** Schematic of the PLOOH biosynthesis pathway. Genes identified from the CRISPR screen were marked in the indicated colors to highlight their contribution to ferroptosis.

To validate the screening results, we treated liver cancer cells with MI-2, EX 527 (selisistat), or AI-10-49. Huh7 and SK-Hep1 cells demonstrated resistance to high-concentration treatment of EX 527 (selisistat) (Extended Data Fig. 2a). Cells treated with AI-10-49 could be partially rescued by ferrostatin-1, however, this effect was observed only within a narrow concentration range (Extended Data Fig. 2b). Cell death induced by MI-2 in Huh7, SK-Hep1, and SNU398 cells could be effectively rescued by ferrostatin-1. However, other types of cell death inhibitors were ineffective in suppressing the cell death induced by MI-2 (Fig. 1e and Extended Data Fig. 2c). We also noted an increase in lipid reactive oxygen species (ROS) levels and mitochondrial damage in Huh7, SK-Hep1 and SNU398 cells treated with MI-2 (Fig. 1f, g and Extended Data Fig. 2d, e). Additionally, MI-2 significantly suppress growth of Huh7 xenografts (Fig. 1h). Treatment with MI-2 led to increased lipid peroxidation levels evidenced by 4-HNE staining, and decreased proliferation, as indicated by Ki67 staining (Fig. 1i).

Here, our findings suggest that MI-2 could represent a novel ferroptosis inducer with potential applications in the treatment of liver cancer. In order to uncover the regulatory network responsible for the sensitivity of MI-2 treatment, we conducted an unbiased genome-wide CRISPR screening (Fig. 1j). This analysis identified the essential role of three genes: ACSL4 (Acyl-CoA synthetase long-chain family member 4), LPCAT3 (lysophosphatidylcholine acyltransferase 3), and AGPAT3 (1-acylglycerol-3-phosphate O-acyltransferase 3), in MI-2-induced cell death (Fig. 1k). ACSL4 is responsible for converting polyunsaturated fatty acids (PUFAs) into their respective acyl-CoAs and plays a crucial role in maintaining PUFA-containing membrane phospholipids (PUFA-PLs)^22^. LPCAT3 and AGPAT3 demonstrate a specific preference for PUFA-CoA as their substrate^5,23,24^. By increasing the levels of PUFA-PLs or PUFA-containing ether phospholipids (PUFA-ePLs), they enhance the susceptibility of cancer cells to ferroptosis (Fig. 1l)^25^. This unbiased genome-wide CRISPR screening further confirms that MI-2-induced cell death primarily occurs through ferroptosis.

### MI-2 directly binds and inhibits the activity of GPX4

Having confirmed the ability of MI-2 to induce ferroptosis in liver cancer cells, we aimed to investigate the underlying mechanism. Ferroptosis inducers can be categorized into two classes: class I ferroptosis inducers act by inhibiting SLC7A11-mediated cystine uptake, leading to reduced levels of intracellular cysteine and glutathione (GSH), as exemplified by erastin; class II ferroptosis inducers, such as RSL3, ML162 and ML210, operate by targeting the enzymatic activity of GPX4^3^. To unravel the mechanism of MI-2 induced ferroptosis, we examined its impact on glutamate release and GSH level. In comparison to erastin, MI-2 demonstrated minimal inhibition of glutamate release (Fig. 2a). Additionally, treatment of Huh7 and SK-Hep1 cells with MI-2, did not lead to a decrease in intracellular GSH content compared to cells treated with erastin (Fig. 2b). These findings suggest that MI-2 induces ferroptosis through a distinct mechanism from class I ferroptosis inducers.

**Figure 2.**
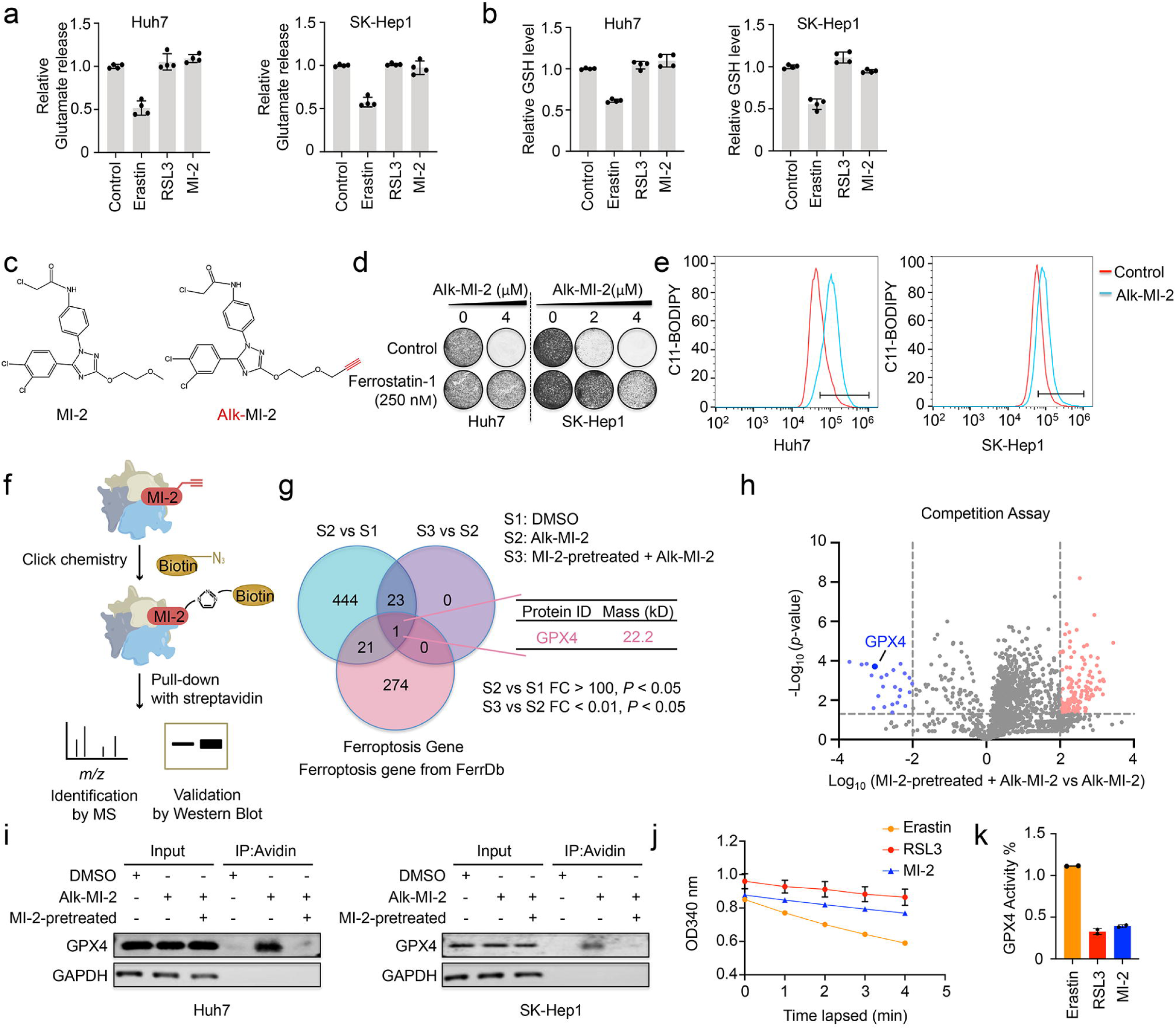
MI-2 directly binds to GPX4 and inhibits the activity of GPX4. **a,** Glutamate release levels of Huh7 and SK-Hep1 cells treated with erastin (system x^−^ inhibitor, 5 μM), RSL3 (GPX4 inhibitor, 125 nM) or MI-2 (1 μM). **b,** GSH levels of Huh7 and SK-Hep1 cells under treatment of erastin (5 μM), RSL3 (250 nM) or MI-2 (2 μM). **c,** The chemical structures of MI-2 and alkyne modified-MI-2, referred to as Alk-MI-2. In Alk-MI-2, the alkyne group is highlighted in red. **d,** Colony formation assays were performed in the presence of Alk-MI-2 or a combination of Alk-MI-2 and ferrostatin-1 for 10-14 days. **e,** Lipid ROS level of Alk-MI-2 treated-cells (Huh7 and SK-Hep1) was assessed through C11-BODIPY fluorescence. **f,** Schematic representation of the activity-based protein profiling (ABPP) performed in SK-Hep1 cells. The target proteins were covalently labeled with Alk-MI-2 probes, followed by their conjugation to an affinity tag (Biotin-N3) via click chemistry. The potential target proteins were identified by mass spectrometry and further validated by western blot. **g,** GPX4 was identified as the only ferroptosis-related target directly bound by MI-2. S1, DMSO treated; S2, Alk-MI-2 treated; S3, Pretreated with MI-2 followed by Alk-MI-2 treatment. **h,** Volcano plot indicated that pretreatment with MI-2 inhibited the binding of GPX4 to Alk-MI-2. **i,** The results of competition assay were validated using western blot in Huh7 and SK-Hep1 cells, which were pretreated with MI-2 followed by Alk-MI-2 treatment. **j,** Time lapsed reduction of NADPH availability in GPX4 assays reaction measured by OD340. **k,** Calculated GPX4 activity followed treatment of erastin, RSL3 and MI-2.

Next, we utilized activity-based protein profiling (ABPP) to identify the primary target of MI-2. Initially, we introduced an alkyne group to modify MI-2 (Fig. 2c) and confirmed that this modification still maintains the capacity of the compound to induce ferroptosis (Fig. 2d, e). The ABPP approach involved labeling target proteins with the Alk-MI-2 probe, followed by biotin conjugation through click chemistry for further enrichment. Subsequently, the candidate target proteins were quantified through mass spectrometry (Fig. 2f). Our analysis identified a total of 489 proteins that could interact with Alk-MI-2. Through a competition assay involving pre-treatment with MI-2, we identified 24 proteins that specifically bind to MI-2. Remarkably, among these 24 candidates, GPX4 emerged as the sole target associated with ferroptosis (Fig. 2g, h). To further confirm the direct binding of MI-2 to GPX4, competition experiments was conducted using western blot assay. As expected, MI-2 pretreatment significantly blocked the direct binding between GPX4 and Alk-MI-2 (Fig. 2i). Then, we aimed to investigate whether the binding of MI-2 to GPX4 could affect GPX4 activity. We performed a GPX4 assay based on the measurement the time-dependent reduction in NADPH availability. Our results showed that both MI-2 and RSL3 were able to suppress GPX4 activity, evidenced by slow reduction rate of NADPH. However, erastin did not exhibit any significant effect on GPX4 activity (Fig. 2j, k).

To compare the response of liver cancer cells to MI-2 and traditional GPX4 inhibitors, we treated a panel cell lines with increasing concentrations of MI-2, RSL3 or ML210 for 10–14 days in colony formation assays. While the response to these compounds varied across cell lines, the effects of MI-2 and GPX4 inhibitors on the panel of liver cancer cell lines were remarkably similar (Extended Data Fig. 3a). It is known that therapy-resistant cancer cells often exhibit altered metabolic states that make them susceptible to ferroptosis induction^5,26^. To investigate this phenomenon, we established lenvatinib-resistant and BLU554-resistant Hep3B cells (Extended Data Fig. 3b) and treated them with RSL3 or MI-2 in colony formation assays. Interestingly, we found that the resistant cells displayed increased sensitivity to both MI-2 and RSL3 treatments, which could be rescued by the addition of ferrostatin-1 (Extended Data Fig. 3c, d). Together, these results strongly indicate that MI-2-induced ferroptosis operates through a mechanism similar to that of traditional class II ferroptosis inducers.

### MI-2 downregulates GPX4 through ubiquitination-mediated degradation

MI-2 is originally identified as an inhibitor of MALT1 protease, which has shown notable efficacy against activated B cell-like diffuse large B cell lymphoma (ABC-DLBCL) in both *in vitro* and *in vivo* studies^27^. In a genome-wide fluorescence-activated cell sorting (FACS)-based CRISPR screening, we unexpected identified MALT1 as one of the most significant hits regulating GPX4 level (Fig. 3a, b). As expected, GPX4 gRNAs were highly enriched in the GPX4^low^ cell population, underscoring the reliability of the genetic screening (Fig. 3b). Additionally, FBXW7, PCBP1 and BAP1 were identified as regulators of GPX4 expression, consistent with their known involvement in ferroptosis regulation (Fig. 3b) ^28–31^. To verify the screening findings, we generated MALT1-knockout PLC/PRF/5 cells and evaluated the levels of GPX4 protein. Compared to parental PLC/PRF/5 cells, MALT1-knockout cells exhibited comparable mRNA but reduced protein level of GPX4 (Fig. 3c, d).

**Figure 3.**
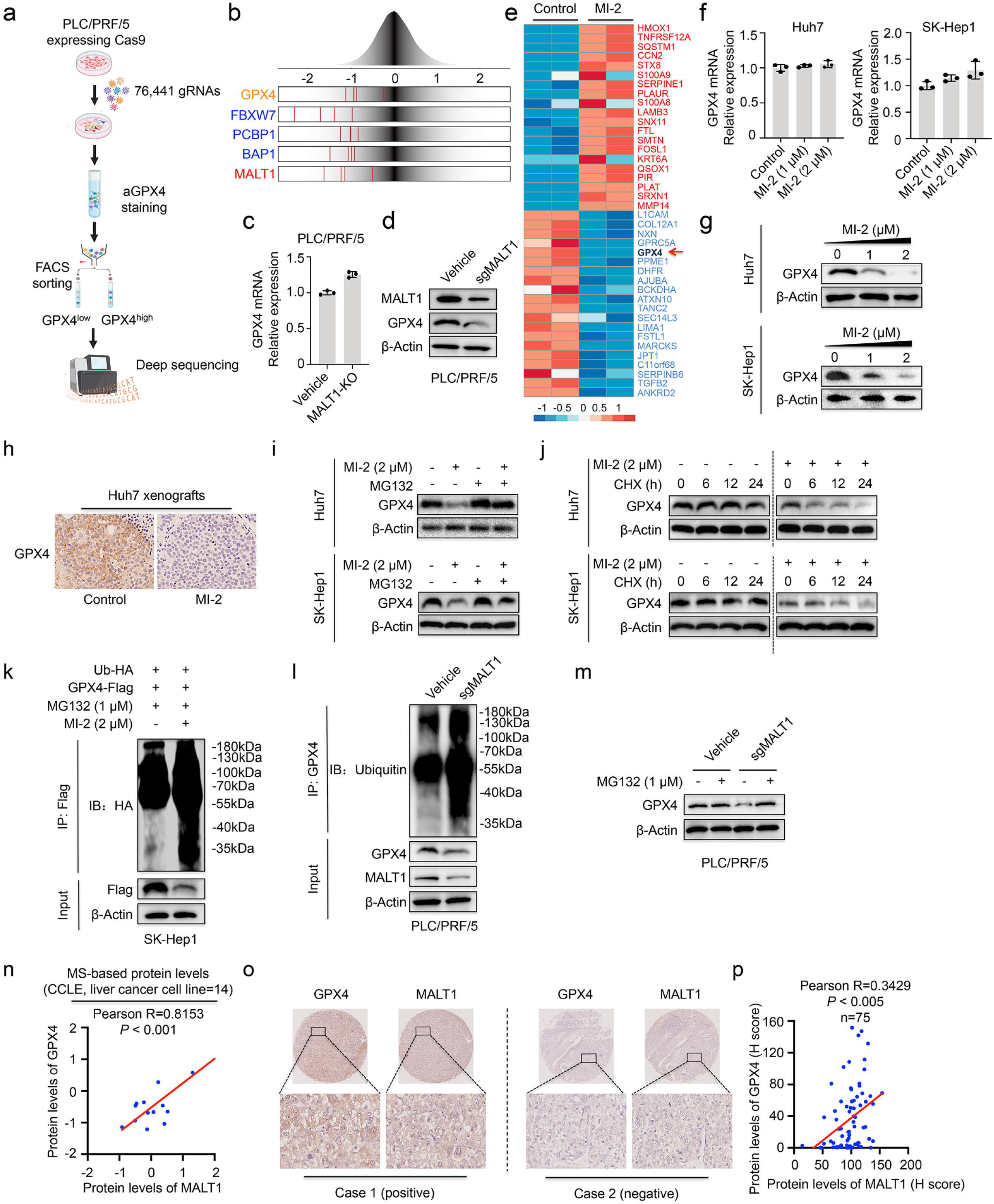
MI-2 facilitates ubiquitous degradation of GPX4. **a,** Schematic illustration of the flow cytometry-based CRISPR-Cas9 screen for modulators of GPX4 expression. **b,** Frequency histograms depicting the log_2_-fold changes of MALT1 gRNAs and GPX4 gRNAs (positive control) in FACS-based genome-wide screening of GPX4 expression in PLC/PRF/5 cells. **c-d,** The expression of GPX4 in MALT1-knockout PLC/PRF/5 cells was measured by qRT-PCR or western blot. **e,** Quantitative proteomics analysis of SK-Hep1 cells in the presence or absence of MI-2 (2 μM) for 4 days. The red arrow indicates the downregulation of GPX4 upon MI-2 treatment. **f-g,** qRT-PCR analyses and western blot were performed to assess the level of GPX4 mRNA and protein in Huh7 and SK-Hep1 cells treated with MI-2 for 48 hours. **h,** Representative images of GPX4 staining performed on formalin-fixed paraffin-embedded Huh7 xenografts from BALB/c nude mice following vehicle or MI-2 (20Lmg/kg) treatment for 12 days. **i,** Western blot analysis of GPX4 in Huh7 and SK-Hep1 cells exposed to MG132 (500 nM), MI-2 (2 μM) or the combination of both for 24 hours. MG132, the inhibitor of proteasome. **j,** Western blot was used to assess the levels of GPX4 proetin in Huh7 and SK-Hep1 cells upon CHX (50 ug/ml) treatment in the presence or absence of MI-2 (2 μM). CHX, the protein synthesis inhibitor. **k,** Immunoprecipitation analysis was performed to assess the ubiquitination level of GPX4 in SK-Hep1 cells transfected with Flag-tagged GPX4 expression plasmids and ub-HA plasmids. **l,** Analysis of GPX4 ubiquitination in MALT1-knockout PLC/PRF/5 cells. **m,** The level of GPX4 protein was assessed by western blot in control or MALT1-knockout PLC/PRF/5 cells treated with or without MG132 (1 μM). **n,** Correlation between the protein levels of MALT1 and GPX4 in liver cancer cell lines, excluding JHH1. **o-p,** Immunohistochemical staining was conducted on liver cancer tissue samples to assess the protein levels of GPX4 and MALT1, and their correlation was subsequently calculated.

Furthermore, quantitative proteomics analysis revealed that MI-2 treatment led to a reduction of GPX4 protein levels in SK-Hep1 cells, which was confirm through *in vitro* and *in vivo* validation (Fig. 3e-h). To extend our investigation, we used another MALT1 inhibitor safimaltib, and observed similar effects on downregulating the protein level of GPX4, without affecting its mRNA expression (Extended Data Fig. 4a, b).

Since MALT1 depletion or inhibition did not affect the RNA transcript levels of GPX4 (Fig. 3c, f), while significantly reducing the GPX4 protein levels (Fig. 3d, g), we hypothesized that the regulation may occur at a posttranscriptional level. To explore the mechanisms involved in MALT1-mediated GPX4 regulation, we treated cells with proteasome or protein synthesis inhibitors and assessed GPX4 stability. We observed that the reduction in GPX4 protein levels induced by MI-2 could be blocked by the proteasome inhibitor MG132 and facilitated by the protein synthesis inhibitor cycloheximide (Fig. 3i, j). Additionally, we noticed increased levels of GPX4 ubiquitination in liver cancer cells upon pharmacological inhibition of MALT1 using MI-2 (Fig. 3k). Consistent with MI-2 treatment, MALT1-knockout cells exhibited increased ubiquitination of GPX4 and the downregulation of GPX4 induced by MALT1 deficiency could be efficiently rescued by the proteasome inhibitor MG132 (Fig. 3l, m).

Through bioinformatics analysis, positive correlations were observed between MALT1 protein levels and GPX4 protein levels in CCLE pan-cancer cell lines, particularly in liver cancer cell lines (Fig. 3n and Extended Data Fig. 4c). However, no substantial correlation was observed regarding the mRNA levels of these two genes (Extended Data Fig. 4c, d). Tissue array analysis of MALT1 and GPX4 in 76 human HCC tissues showed a significantly positive correlation between their protein levels (Fig. 3o, p), further suggesting a potential positive regulatory relationship between MALT1 and GPX4.

In summary, these findings establish MALT1 as a regulator of GPX4 protein levels. Besides direct affecting GPX4 activity, MI-2 exerts a dual modulatory effect on GPX4 stability at the posttranscriptional level.

### GPX4 inhibition is synergistic with sorafenib in liver cancer cells

Given that many liver cancer cell lines demonstrate insensitivity to MI-2, we aimed to investigate whether combination treatment could enhance the susceptibility of these cells to ferroptosis induction. Initially, we confirmed that MI-2 effectively reduced the protein levels of GPX4 in Hep3B, SNU449 and MHCC97H cells, irrespective of their sensitivity to ferroptosis induction (Fig. 4a). To determine if conventional therapeutic drugs used in HCC treatment could synergize with MI-2 in these ferroptosis-insensitive cells, we treated Hep3B, SNU449 and MHCC97H cells with MI-2 and sorafenib, regorafenib, lenvatinib or the indicated combinations. Strong synergies between MI-2 with sorafenib or regorafenib were observed in Hep3B, SNU449 and MHCC97H cells (Fig. 4b and Extended Data Fig. 5a). However, there is no synergistic effect when cells were treated with MI-2 and lenvatinib (Extended Data Fig. 5a). Sorafenib, regorafenib and lenvatinib are all clinically approved multi-kinase inhibitors used in the treatment of advanced HCC. Regorafenib is structurally similar to sorafenib^9,32,33^, which can explain the similar synergic effects observed when combining either sorafenib or regorafenib with MI-2. Comparable results were obtained when combining RSL3 with sorafenib or regorafenib (Extended Data Fig. 5b). Besides, we noted ferroptosis-specific mitochondrial alterations and increasing lipid peroxides accumulation in liver cancer cell lines treated with sorafenib when combined with MI-2 (Fig. 4c, d) or RSL3 (Extended Data Fig. 5c, d).

**Figure 4.**
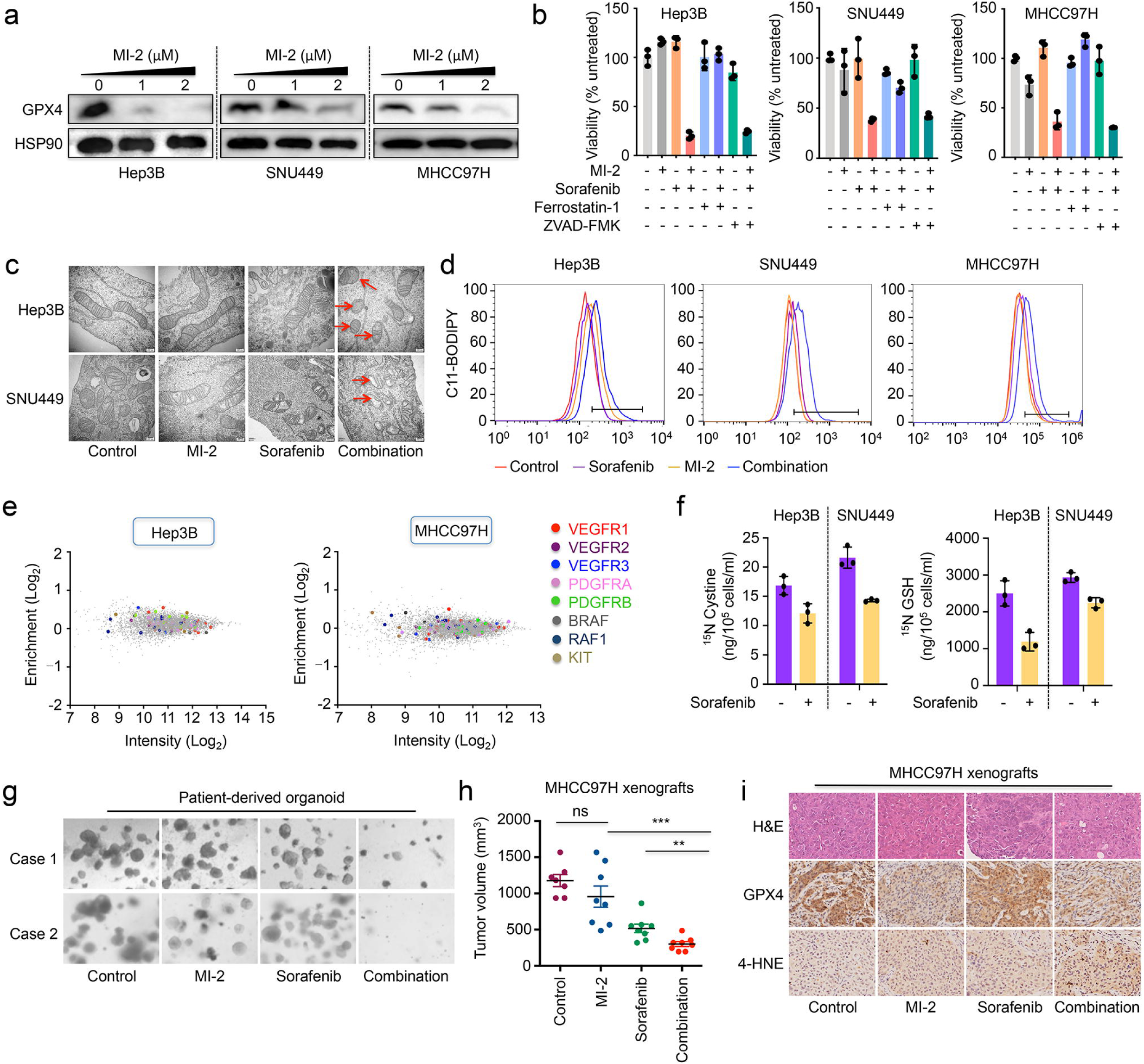
MI-2 synergies with sorafenib in liver cancer cells. **a,** Western blot analysis of GPX4 in three HCC cell lines (insensitive to ferroptosis induction: Hep3B, SNU449 and MHCC97H) treated with MI-2 for 3-4 days. **b,** Quantification of CellTiter-Blue viability assays of Hep3B, SNU449 and MHCC97H cells exposed to MI-2 (0.25-2 μM), sorafenib (2.5 μM), ferrostatin-1 (250 nM), ZVAD-FMK (1.25 μM) or the indicated combination. **c,** Hep3B and SNU449 cells were treated with MI-2, sorafenib or the combination for 3 days and ultrastructural analysis was performed using transmission electron microscopy. Red arrows indicate damaged mitochondria. **d,** Lipid ROS level was assessed by C11-BODIPY fluorescence on the Hep3B, SNU449 and MHCC97H cells treated with sorafenib, MI-2 or the combination. **e,** Representation of the relative abundance of the gRNA barcode sequences from the kinome screens. Common targets of sorafenib were not identified as high-confidence synthetic lethal genes with GPX4 inhibition in Hep3B and MHCC97H cells. **f,** ^15^N labeled L-cysteine was add to the cell culture medium. Cystine uptake (^15^N cystine and ^15^N GSH level) was accessed in Hep3B and SNU449 cells upon sorafenib (10 μM) treatment for 3 days. **g,** Representative images showed the response of patient-derived organoids (PDOs) to MI-2 (1 μM), sorafenib (2.5 μM) or the combination of both. **h,** Tumor volumes of MHCC97H xenografts in BALB/c nude mice following vehicle, MI-2 (20 mg/kg), sorafenib (30 mg/kg) or combination treatment for 14 days. \*\**P* < 0.01, \*\*\**P*<0.001. **i,** Representative images of H&E, GPX4 and 4-HNE staining performed on formalin-fixed paraffin-embedded MHCC97H xenografts.

Through kinome-based CRISPR screenings, we found that loss of kinase targets of sorafenib did not synergize with GPX4 inhibitor RSL3, indicating the synergy between GPX4 inhibitor and sorafenib may be independent of its common targets (Fig. 4e). As a multi-kinase inhibitor, sorafenib has been shown to induce ferroptosis by targeting SLC7A11^21,34^. We next used ^15^N labeled L-cysteine to trace cystine uptake and biosynthesis of GSH. As expected, sorafenib treatment induced a notable reduction in both cystine uptake and biosynthesis of GSH, as evidenced by the utilization of ^15^N-labeled L-cysteine (Fig. 4f). SLC7A11, as one of the subunits of system x_c_ ^−^, plays a crucial role in cystine uptake^35^. We found that expression of SLC7A11 was significantly associated with the sensitivity to GPX4 inhibitor in panels of liver cancer cell line and pan-cancer cell lines (Extended Data Fig. 5e). Together, these findings suggest that sorafenib may synergize with MI-2 through its activity against SLC7A11.

In order to enhance the clinical relevance of our findings, we evaluated synergistic effect of this combination in patient-derived liver cancer organoid (PDO) models. The combination of MI-2 and sorafenib notably suppressed proliferation of the organoids (Fig. 4g). To assess whether these *in vitro* findings could be recapitulated *in vivo*, we established subcutaneous tumors in nude mice using MHCC97H cells. Consistent with *in vitro* results, the combined administration of MI-2 and sorafenib demonstrated a substantial inhibition of tumor growth, accompanied by decreased GPX4 expression and increased lipid peroxidation levels in tumor tissues (Fig. 4h, i).

### MI-2 is synergistic with sorafenib or regorafenib in multiple caner models

Sorafenib has been approved by the FDA for the treatment of thyroid cancer and kidney cancer, while regorafenib has been approved for colorectal cancer treatment^12,13,36,37^. The remarkable synergy observed between MI-2 and sorafenib in liver cancer cells have inspired us to investigate the potential of this combination therapy in treating these three types of cancer. Encouragingly, synergistic effects on cell proliferation were observed when sorafenib or regorafenib was combined with MI-2 in thyroid cancer, kidney cancer and colorectal cancer cell lines (Extended Data Fig. 4a-c). CellTiter-Blue viability assay confirmed that the cell death induced by these combinations was attributable to ferroptosis, as evidenced by the rescue effects of ferrostatin-1 (Extended Data Fig. 4d-f). Furthermore, an increase in lipid ROS levels was noted in KTC-1, OSRC-2 and LoVo cells upon combination treatment (Extended Data Fig. 6g). Consistent with *in vitro* results, mice bearing OSRC-2 and LoVo xenografts that received combination therapy exhibited a significant reduction in tumor burden compared to those receiving monotherapy (Extended Data Fig. 6h, j). Ferroptosis induction *in vivo* was also evident in tumor tissues with combination treatment, as demonstrated by GPX4 and 4-HNE staining (Extended Data Fig. 6i, k). Collectively, these findings suggest that combining sorafenib or regorafenib with GPX4 inhibition using MI-2 could represent a broadly applicable strategy for a diverse range of malignancies.

## Discussion

Inhibition of GPX4 with small molecules has proven to be challenging due to the flat surface surrounding the active site and the absence of known regulatory sites^7^. The most widely used inhibitor is RSL3, but it lacks favorable drug-like properties, limiting its clinical development. Other inhibitors including ML162 and ML210 also show potent ferroptosis-inducing capabilities but face similar drawbacks^7,8^. In order to identify novel and effective ferroptosis inducer, a compound screening approach was employed, and MI-2 was identified as a strong inducer of ferroptosis, which was initially recognized as an inhibitor of MALT1 protease. Subsequent unbiased genome-wide CRISPR screening further confirmed that cell death induced by MI-2 predominantly occurs through ferroptosis. Currently, ferroptosis inducers can be primarily categorized into two classes based on their mechanisms of action: those that inhibit SLC7A11-mediated cystine uptake and those that block the enzymatic activity of GPX4. In our study, activity-based protein profiling (ABPP) assays yielded compelling evidence of a direct interaction between MI-2 and GPX4. This interaction led to a notable decrease in the activity of GPX4, suggesting that MI-2 can be classified as a class II ferroptosis inducer. These findings demonstrate the specific effects of MI-2 on targeting GPX4 and hold great promise as a starting point for the development of drug-like GPX4 inhibitors with translational potential.

Recent research has highlighted that GPX4 can undergo various post-translational modifications (PTMs), including ubiquitination, phosphorylation and glycosylation^38^. These PTMs affect the protein level and activity of GPX4, suggesting that targeting these processes could be a potential therapeutic approach for ferroptosis-related diseases. Studies have unveiled specific mechanisms of GPX4 regulation, such as TRIM25-mediated K48-linked ubiquitination triggering its proteasomal degradation^39^, and OTUD5 conferring ferroptosis resistance by stabilizing GPX4^40,41^. Building upon these insights, our study employed a FACS-based genome-wide genetic screen to systematically identify modulators of GPX4. Unexpectedly, we discovered that MALT1, previously recognized as the target of MI-2, emerged as one of the most significant regulators of GPX4 levels. Lorena Fontan et al. developed MI-2, which directly binds to MALT1 and suppresses its protease function, thereby inhibiting ABC-DLBCL both *in vitro* and *in vivo*^27^. Notably, MI-2 displayed low toxicity in mice while effectively reducing the growth of Huh7 xenografts, accompanied by increased levels of lipid peroxidation in tumor tissues. A recent study suggested that MI-2 could sensitize cancer cells to glutathione (GSH) depletion by efficiently inhibiting multiple deubiquitinating enzymes (DUBs)^42^. Consistent with this, our study confirmed that MI-2 promotes the degradation of GPX4, aligning with the effects of MI-2 on DUBs. However, further investigations are required to elucidate the specific DUBs involved in the link between MALT1 and GPX4 degradation.

Combining different agents with ferroptosis inducers has shown promise in enhancing the susceptibility of cancer cells to ferroptosis induction^17,43,44^. In our study, we utilized sorafenib or regorafenib as combination agents to induce ferroptosis. Both sorafenib and regorafenib exhibited synergistic effects when combined with MI-2 or RSL3 in inducing ferroptosis. Additionally, both our findings and other studies have indicated that sorafenib also exerts activity against SLC7A11, a key component involved in ferroptosis regulation^20^. Our kinome screening data further demonstrated that the synergy between sorafenib and GPX4 inhibitors is independent of their kinase targets. This synergy can be attributed to the co-targeting of SLC7A11 and GPX4, key components of system X_c_^-^ - GSH -GPX4 axis, which helps overcome the intrinsic resistance of cancer cells to ferroptosis induction. Considering that sorafenib or regorafenib has been approved for use in various cancer types, including advanced liver cancer, thyroid cancer, kidney cancer, and colorectal cancer, this combination strategy holds potential for further investigation.

An important limitation of our study is the lack of detailed structure-activity relationship analysis between MI-2 and GPX4. While we have demonstrated that MI-2 directly interacts with GPX4 and inhibits its activity, further experimental evidence, like cryo-electron microscopy reconstructions, is required to gain a deeper understanding of this interaction. Additionally, although we have conducted genome-wide screening, quantitative proteomics analysis, and ubiquitination assays, which support the involvement of MALT1 in the ubiquitination-mediated degradation of GPX4 induced by MI-2, it is possible that other downstream proteins also play a role in the MALT1-dependent regulation of ferroptosis. Future investigations should focus on elucidating these aspects.

In summary, we have discovered a small molecule known as MI-2 that effectively inhibits the activity of GPX4 by directly interacting with it. Moreover, MI-2 induces the degradation of GPX4, which relies on its commonly recognized target MALT1. To our knowledge, MI-2 is a novel compound with dual modulatory effects on both the activity and stability of GPX4. In multiple cancer models, MI-2 has shown synergistic effects in inducing ferroptosis when combined with sorafenib or regorafenib. We anticipate that this strategy should undergo further preclinical evaluation to determine its potential for clinical translation.

## Methods

### Compound screens

Huh7 and SK-Hep1 were screened with a compound library (n=2103, 2 μM) in the presence or absence of 250 nM ferrostatin-1 for 4 days in two replicates. Cell viability was assessed by CellTiter-Glo. Compounds exhibiting an impact on cell viability that could be rescued by ferrostatin-1 (at least 1.5-fold increase in the group treated with combination of the compound and ferrostatin-1 compared to the group treated with the compound alone) were selected for further investigation.

### Lipid peroxidation assessment

Briefly, 200,000-250,000 cells per well were seeded in 6-well plates and then cells were treated with indicated compounds for 3-4 days. Cells were harvested and resuspended in PBS containing C11-BODIPY (581/591) (2 μM) (D3861, Invitrogen) at 37°C for 30 min. Subsequently, cells were analyzed using a flow cytometer equipped with 488 nm laser for excitation. A minimum of 10,000 cells were analyzed per sample.

### Xenografts

Huh7 cells (1 × 10^7^ cells per mouse) were injected subcutaneously into the right posterior flanks of 6-week-old BALB/c nude mice (male, 6 mice per group). Tumor volume based on caliper measurements was calculated using modified ellipsoidal formula: tumor volume = 1/2 length × width^2^. After tumor establishment, the mice were randomly assigned to 6 days / week treatment with vehicle, or MI-2 (20mg/kg, intraperitoneal injection). For combination treatment assay, MHCC97H, OSRC-2 and LoVo cells (1 × 10^7^ cells per mouse) were injected subcutaneously into the right posterior flanks of 6-week-old BALB/c nude mice (male, 6-8 mice per group). Mice were randomly assigned to different treatment groups: vehicle, MI-2 (20mg/kg, intraperitoneal injection, 6 days per week), sorafenib (30mg/kg, oral gavage, 3 or 6 days per week), regorafenib (5mg/kg, oral gavage, 6 days per week) or a drug combination in which each compound was administered at the same dose and schedule as the single agent.

### CRISPR-Cas9 genetic screens

To identify genes whose knockout may confer resistance to MI-2, CRISPR-Cas9 genetic screens were conducted in SK-Hep1 cells. The genome-wide CRISPR library was introduced into SK-Hep1 cells by lentiviral transduction. The T0 arm was used as one of the controls after puromycin selection. The cells were then divided into two groups with or without treatment of MI-2 (700 nM) for 45 days. The alterations in library representation were evaluated through Illumina deep sequencing to determine changes in gRNA abundance.

### FACS-based CRISPR-Cas9 screens

Fluorescence-activated cell sorting (FACS)-based genome-wide CRISPR screens were performed using Brunello lentiviral pooled libraries in PLC/PRF/5 cells. The cells selected by puromycin were first stained with LIVE/DEAD™ Fixable Near-IR Dead Cell Stain (L10119, Invitrogen) to exclude nonviable cells, and then fixed and permeabilized with Fixation / Permeabilization solution (88-8824-00, Thermo Fisher Scientific). After washing, the cells were incubated with the anti-GPX4 antibody for 45 minutes at 4°C, followed by incubation with the appropriate dilutions of Alexa Fluor secondary antibody for 30 minutes at 4°C in the dark. The cells were then washed with FACS buffer, and the top 5% (cells with the highest signals) and bottom 5% (cells with the lowest signals) populations were sorted and collected for deep sequencing and bioinformatic analysis.

### Ubiquitination assay

A co-transfection of HA-tagged ubiquitin plasmid with Flag-tagged GPX4 was conducted to evaluate the ubiquitination levels of Flag-GPX4 in cells. The transfected cells were treated with or without MI-2 in the presence of proteasome inhibitor MG132 . Subsequently, the cells were lysed in cold Pierce IP Lysis buffer (87787, Thermo Fisher Scientific) and cleared by centrifugation. An appropriate amount of protein was then incubated with anti-Flag beads overnight at a constant rotation. The beads were washed three times with IP lysis buffer, followed by immunoblotting analysis with anti-HA antibody. To detect the ubiquitination levels of endogenous GPX4 in parental and MALT1 knockout PLC/PRF/5 cells, we lysed the cells and subjected them to immunoprecipitation with anti-GPX4 antibody and protein G-conjugated magnetic beads overnight at constant rotation, followed by immunoblotting analysis with anti-ubiquitin antibody.

### Generation of liver cancer organoid

Liver cancer tissues were minced and digested with PBS supplemented with collagenase type IV at 37°C for 30 to 60 minutes. The suspension was filtered through a 100-μm cell strainer and then centrifuged. The pellet was resuspended in cold organoid culture medium and then combined with Matrigel at a 1:2 ratio to achieve a cell density of 4000 cells per 50 μL before being seeded into a 12-well culture plate. Organoid culture medium was added to each well after solidification and organoids were cultured in a humidified incubator at 37°C with 5% CO_2_. The organoids were then cultured for 5-7 days in medium supplemented with the specific drugs. Image acquisition was by means of ZESS AxioObserver7.

### Labeling of live cells with Alk-MI-2 and streptavidin pulldown

SK-Hep1 cells were treated with DMSO or Alk-MI-2 (5 μM) for 6 hours or pre-treated with MI-2 (10 μM) for 3 hours followed by incubation with Alk-MI-2 (5 μM) for an additional 6 hours. After washing with PBS buffer, cells were lysed in RIPA Buffer (50 mM Tris-HCl, pH 7.4, 150 mM NaCl, 1% Triton X-100, 1% deoxycholate, 0.1% SDS, 10% glycerol, and protease inhibitors) for click chemistry to conjugate with Biotin-N3. Biotin-N3 (final concentration 200 μM), TBTA (final concentration 600 μM), CuSO4 (final concentration 1 mM), and TCEP (final concentration 1 mM) were added to samples in order, and reaction mixture was incubated at room temperature for 1 hours. After click chemistry, proteins (2 mg) were precipitated using methanol / chloroform and resolubilized in 8M Urea buffer (8M Urea, 50 mM Tris-HCl, pH 7.4, 150 mM NaCl, and protease inhibitors). After dilution in Tris-HCl buffer (50 mM Tris-HCl, pH 7.4, 150 mM NaCl, and protease inhibitors) to reduce the Urea concentration below 2M, samples were incubated with streptavidin beads (20349, Thermo) at room temperature for 4 hours with rotation. The bound proteins were eluted with protein loading buffer and resolved by SDS-PAGE or digested by trypsin for mass spectrometric analysis.

### Mass spectrometric analysis of streptavidin pulldown

#### On-beads digestion

Beads were suspended in 30 μL of 8 M Urea in PBS and treated with10mM DTT for 30 min at 55℃ followed by the treatment of 25 mM iodoacetamide for 20 min at room temperature (keep in dark). The supernatant was collected, and the urea concentration was diluted to 2 M with 50mM NH_4_HCO_3_. Then the solution was treated with 0.2 μg trypsin for digestion at 37°C overnight. Digested peptides were enriched by C18 tips for LC-MS/MS detection.

#### LC-MS/MS Analysis

The peptide mixtures were analyzed using an on-line EASY-nL-LC 1000 coupled with an Orbitrap Fusion mass spectrometer. The sample was loaded directly onto a 15-cm home-made capillary column (C18-AQ, 1.9 mm, Dr. Maisch, 100 mm I.D.) at a flow rate of 300 nl/min. Mobile phase A consisted of 0.1% formic acid, 2% acetonitrile and 98% LC-MS grade H_2_O and mobile phase B consisted of 0.1% formic acid, 2% LC-MS grade H_2_O and 98% acetonitrile. For IP-MS analysis, data were acquired in a data-dependent mode with one full MS1 scan in the Orbitrap (m/z: 200–1800; resolution: 120 000; AGC target value: 400 000 and maximal injection time: 50 ms), followed by MS2 scan in the in the Orbitrap (32% normalized collision energy; resolution: 30 000; AGC target value: 10 000; maximal injection time: 100 ms).

#### MS Data Analysis

MS/MS raw spectra were processed using MaxQuant software (version 1.5.3.30). The human protein sequence database containing 20410 sequence entries in the Swiss-Prot database downloaded on October 12, 2018 was used for database search. Trypsin was set as the enzyme, and the maximum missed cleavage was set to 2. The first-search peptide mass tolerance and main-search peptide tolerance were set to 20 and 4.5 ppm. The MS/MS match tolerance was set to 0.5 Da for ITMS and 20 ppm for FTMS. A fixed carbamidomethyl modification of cysteine and variable modifications including oxidation on methionine were set. “Match between runs” was applied, and the match time window was set to within 2 min. The false discovery rate (FDR) was controlled with a decoy database and set to no more than 1%. For analysis, protein hits are required to be detected in repeat samples. Undetected MS intensities are set as 10000 for calculation.

## Acknowledgments

This work was funded by grants from National Natural Science Foundation of China (82330095, 81920108025, 82072633, 82122047, 82203044, 82202062 and 82003171), Shanghai Pilot Program for Basic Research - Shanghai Jiao Tong University (21TQ1400225), Innovative research team of high-level local universities in Shanghai, Central University Outstanding Youth Team Cultivation Program, National Facility for Translational Medicine (TMSK-2021-129), Shanghai Municipal Education Commission-Gaofeng Clinical Medicine Grant Support (20181703).

## Author contributions

C.W., C.S., W.Q. and R.B. supervised all of the research. J.W., C.W. and W.Q. wrote the manuscript. J.W. and L.L. designed, performed and analyzed *in vitro* and *in vivo* experiments. Q.Z. and J.Z. performed compound screeing. B.Y. performed activity-based protein profiling assays. B.M., A.F. and C.S. performed FACS-based genome-wide screening. S.J. provided technical support for PDO models. G.J. provided clinical samples and performed the tissue microarray and scoring; B.L., J.H., S.W. and X.M. provided support for the project. All authors commented on the manuscript.

## References

1 Dixon, S. J. et al. Ferroptosis: an iron-dependent form of nonapoptotic cell death. Cell 149, 1060–1072 (2012).

2 Stockwell, B. R. et al. Ferroptosis: A Regulated Cell Death Nexus Linking Metabolism, Redox Biology, and Disease. Cell 171, 273–285 (2017).

3 Yang, W. S. et al. Regulation of ferroptotic cancer cell death by GPX4. Cell 156, 317–331 (2014).

4 Lei, G., Zhuang, L. & Gan, B. Targeting ferroptosis as a vulnerability in cancer. Nat Rev Cancer 22, 381–396, doi:10.1038/s41568-022-00459-0 (2022).

5 Viswanathan, V. S. et al. Dependency of a therapy-resistant state of cancer cells on a lipid peroxidase pathway. Nature 547, 453–457 (2017).

6 Wang, M. E. et al. RB1-deficient prostate tumor growth and metastasis are vulnerable to ferroptosis induction via the E2F/ACSL4 axis. J Clin Invest 133 (2023).

7 Liu, H. et al. Small-molecule allosteric inhibitors of GPX4. Cell Chem Biol 29, 1680–1693.e1689 (2022).

8 Eaton, J. K. et al. Selective covalent targeting of GPX4 using masked nitrile-oxide electrophiles. Nat Chem Biol 16, 497–506 (2020).

9 Vogel, A., Meyer, T., Sapisochin, G., Salem, R. & Saborowski, A. Hepatocellular carcinoma. Lancet 400, 1345–1362 (2022).

10 Sung, H., et al. Global Cancer Statistics 2020: GLOBOCAN Estimates of Incidence and Mortality Worldwide for 36 Cancers in 185 Countries. CA Cancer J Clin 71, 209–249, (2021).

11 EASL Clinical Practice Guidelines: Management of hepatocellular carcinoma. J Hepatol 69, 182–236 (2018).

12 Kudo, M. et al. Lenvatinib versus sorafenib in first-line treatment of patients with unresectable hepatocellular carcinoma: a randomised phase 3 non-inferiority trial. Lancet 391, 1163–1173 (2018).

13 Llovet, J. M. et al. Sorafenib in advanced hepatocellular carcinoma. N Engl J Med 359, 378–390 (2008).

14 Cheng, A. L. et al. Updated efficacy and safety data from IMbrave150: Atezolizumab plus bevacizumab vs. sorafenib for unresectable hepatocellular carcinoma. J Hepatol 76, 862–873 (2022).

15 Finn, R. S. et al. Atezolizumab plus Bevacizumab in Unresectable Hepatocellular Carcinoma. N Engl J Med 382, 1894–1905 (2020).

16 Zhu, A. X. et al. Molecular correlates of clinical response and resistance to atezolizumab in combination with bevacizumab in advanced hepatocellular carcinoma. Nat Med 28, 1599–1611 (2022).

17 Zheng, C. et al. Donafenib and GSK-J4 Synergistically Induce Ferroptosis in Liver Cancer by Upregulating HMOX1 Expression. Adv Sci (Weinh) 10, e2206798, (2023).

18 Conche, C. et al. Combining ferroptosis induction with MDSC blockade renders primary tumours and metastases in liver sensitive to immune checkpoint blockade. Gut 72, 1774–1782 (2023).

19 Yao, F. et al. A targetable LIFR-NF-κB-LCN2 axis controls liver tumorigenesis and vulnerability to ferroptosis. Nat Commun 12, 7333 (2021).

20 Yuan, S. et al. Sorafenib attenuates liver fibrosis by triggering hepatic stellate cell ferroptosis via HIF-1α/SLC7A11 pathway. Cell Prolif 55, e13158 (2022).

21 Gao, R. et al. YAP/TAZ and ATF4 drive resistance to Sorafenib in hepatocellular carcinoma by preventing ferroptosis. EMBO Mol Med 13, e14351 (2021).

22 Doll, S. et al. ACSL4 dictates ferroptosis sensitivity by shaping cellular lipid composition. Nat Chem Biol 13, 91–98 (2017).

23 Gupta, G. et al. Exploring ACSL4/LPCAT3/ALOX15 and SLC7A11/GPX4/NFE2L2 as potential targets in ferroptosis-based cancer therapy. Future Med Chem 15, 1209–1212 (2023).

24 Zou, Y. et al. Plasticity of ether lipids promotes ferroptosis susceptibility and evasion. Nature 585, 603–608 (2020).

25 Hadian, K. & Stockwell, B. R. A roadmap to creating ferroptosis-based medicines. Nat Chem Biol 17, 1113–1116 (2021).

26 Hangauer, M. J. et al. Drug-tolerant persister cancer cells are vulnerable to GPX4 inhibition. Nature 551, 247–250 (2017).

27 Fontan, L. et al. MALT1 small molecule inhibitors specifically suppress ABC-DLBCL in vitro and in vivo. Cancer Cell 22, 812–824 (2012).

28 Zhang, Z. et al. RNA-binding protein ZFP36/TTP protects against ferroptosis by regulating autophagy signaling pathway in hepatic stellate cells. Autophagy 16, 1482–1505, (2020).

29 Huang, G. et al. The lncRNA BDNF-AS/WDR5/FBXW7 axis mediates ferroptosis in gastric cancer peritoneal metastasis by regulating VDAC3 ubiquitination. Int J Biol Sci 18, 1415–1433 (2022).

30 Lee, J., You, J. H. & Roh, J. L. Poly(rC)-binding protein 1 represses ferritinophagy-mediated ferroptosis in head and neck cancer. Redox Biol 51, 102276, (2022).

31 Zhang, Y. et al. BAP1 links metabolic regulation of ferroptosis to tumour suppression. Nat Cell Biol 20, 1181–1192 (2018).

32 Bruix, J. et al. Regorafenib for patients with hepatocellular carcinoma who progressed on sorafenib treatment (RESORCE): a randomised, double-blind, placebo-controlled, phase 3 trial. Lancet 389, 56–66 (2017).

33 Carr, B. I. et al. Fluoro-Sorafenib (Regorafenib) effects on hepatoma cells: growth inhibition, quiescence, and recovery. J Cell Physiol 228, 292–297 (2013).

34 Huang, W. et al. ABCC5 facilitates the acquired resistance of sorafenib through the inhibition of SLC7A11-induced ferroptosis in hepatocellular carcinoma. Neoplasia 23, 1227–1239 (2021).

35 Wang, L. et al. ATF3 promotes erastin-induced ferroptosis by suppressing system Xc^-^. Cell Death Differ 27, 662–675 (2020).

36 Brose, M. S. et al. Sorafenib in radioactive iodine-refractory, locally advanced or metastatic differentiated thyroid cancer: a randomised, double-blind, phase 3 trial. Lancet 384, 319–328 (2014).

37 Grothey, A. et al. Regorafenib monotherapy for previously treated metastatic colorectal cancer (CORRECT): an international, multicentre, randomised, placebo-controlled, phase 3 trial. Lancet 381, 303–312 (2013).

38 Wang, Y. et al. Targeting epigenetic and posttranslational modifications regulating ferroptosis for the treatment of diseases. Signal Transduct Target Ther 8, 449, (2023).

39 Li, J. et al. Tumor-specific GPX4 degradation enhances ferroptosis-initiated antitumor immune response in mouse models of pancreatic cancer. Sci Transl Med 15, eadg3049 (2023).

40 Chu, L. K. et al. Autophagy of OTUD5 destabilizes GPX4 to confer ferroptosis-dependent kidney injury. Nat Commun 14, 8393 (2023).

41 Liu, L. et al. Deubiquitinase OTUD5 as a Novel Protector against 4-HNE-Triggered Ferroptosis in Myocardial Ischemia/Reperfusion Injury. Adv Sci (Weinh) 10, e2301852 (2023).

42 Harris, I. S. et al. Deubiquitinases Maintain Protein Homeostasis and Survival of Cancer Cells upon Glutathione Depletion. Cell Metab 29, 1166–1181.e1166 (2019).

43 Schmitt, A. et al. BRD4 inhibition sensitizes diffuse large B-cell lymphoma cells to ferroptosis. Blood 142, 1143–1155 (2023).

44 Rodencal, J. et al. Sensitization of cancer cells to ferroptosis coincident with cell cycle arrest. Cell Chem Biol (2023).

