## Supplementary figure legends for "Identification of a Novel Ferroptosis Inducer with Dual Modulatory Effects on GPX4 Activity and Stability"

**Figure S1. GPX4 is a potential therapeutic target for liver cancer**

**a,** The mRNA level of GPX4 in tumor tissues and non-tumor tissues in the cohort of TCGA and ICGC database. **b,** Typical immunostaining images of GPX4 for HCC patients. **c，**Kaplan-Meier curves indicating that high level of GPX4 mRNA correlates with poor prognosis of patients with HCC from the TCGA cohort (n=365). **P* < 0.05. **d-e,** The level of GPX4 knockdown in liver cancer cell lines was measured by qRT-PCR or western blot. **f,** Cell viability was assessed by colony formation assay based on GPX4 knockdown.

**Figure S2. Effects of MI-2 on ferroptosis induction in liver cancer cells**

**a-b,** Colony formation assays were performed on Huh7 and SK-Hep1 cells treated with EX527 or AI-10-49 combined with ferrostatin-1 (250 nM) for 10–14 days. **c,** Colony formation assays of SNU398 cells treated with MI-2 combined with ferrostatin-1, ZVAD-FMK, necrostatin-1, or 3-Methyladenine, respectively for 10–14 days. **d,** SNU398 cells were treated with MI-2 (2 μM) for 72 hours and lipid ROS level was assessed by C11-BODIPY fluorescence through flow cytometry. **e,** Ultrastructural analysis of MI-2-treated SNU398 cells using transmission electron microscopy. Red arrows indicate damaged mitochondria.

**Figure S3. MI-2 exhibits the characteristics of classical GPX4 inhibitors**

**a,** Long-term colony formation assays were performed on liver cancer cell lines treated with increasing concentrations of MI-2, RSL3 or ML210, respectively. **b,** Colony formation assays were conducted on lenvatinib-resistant and BLU554-resistant Hep3B cells, as well as their respective controls, following treatment with BLU554 or lenvatinib for a duration of 10-14 days. **c-d,** Colony formation assays demonstrate increased sensitivity of lenvatinib-resistant or BLU554-resistant Hep3B cells to MI-2 or RSL3 treatment.

**Figure S4. MALT1 inhibition downregulates protein level of GPX4**

**a,** qRT-PCR analyses were performed to assess the level of GPX4 mRNA in Huh7 and SK-Hep1 cells treated with MALT1 inhibitor safimaltib. **b,** Western blot analysis of GPX4 and MALT1 protein levels in Huh7 and SK-Hep1 cells upon the treatment of safimaltib for 3-4 days. **c,** Correlation between the mRNA or protein levels of MALT1 and GPX4 in pan-cancer cell lines. **d,** Correlation between the mRNA levels of MALT1 and GPX4 in liver cancer cell lines, excluding JHH1.

**Figure S5. GPX4 inhibition synergies with sorafenib in liver cancer cells**

**a-b,** Colony formation assays of Hep3B, SNU449 and MHCC97H cells treated with MI-2 (0.5-1 μM) or RSL3 (125-250 nM) combined with sorafenib, regorafenib and lenvatinib, respectively for 10-14 days. **c,** Hep3B and SNU449 cells were treated with RSL3, sorafenib or the combination for 3 days and ultrastructural analysis was performed using transmission electron microscopy. Red arrows indicate damaged mitochondria. **d,** Lipid ROS level was assessed by C11-BODIPY fluorescence on the Hep3B and SNU449 cells treated with sorafenib (2.5 μM), RSL3 (250 nM) or the combination. **e,** Correlation between the expression of SLC7A11 and the drug sensitivity of RSL3 in liver cancer and pan-cancer cell lines. The x-axis depicts the AUC of RSL3. Lower values on the X-axis imply greater drug sensitivity.

**Figure S6. The combination of MI-2 and sorafenib or regorafenib triggers ferroptosis in multiple types of cancer**

**a-b,** Colony formation assays were performed on thyroid cancer cells (Cal62 and KTC-1) and renal cancer cells (OSRC-2 and ACHN) treated with MI-2 combined with sorafenib for 10-14 days. **c,** Colony formation assays of colorectal cancer cells (LoVo and SW480) exposed to the combination of MI-2 and regorafenib for 10-14 days. **d-f,** CellTiter-Blue viability assays of Cal62, KTC-1, OSRC-2, ACHN, LoVo and SW480 cells exposed to sorafenib or regorafenib, MI-2, ferrostatin-1, ZVAD-FMK or the indicated combinations. **g,** KTC-1, OSRC-2 and LoVo cells were treated with MI-2 combined with sorafenib or regorafenib for 3-5 days and lipid ROS level was assessed by C11-BODIPY fluorescence through flow cytometry. **h,** Tumor volumes of OSRC-2 xenografts in BALB/c nude mice following vehicle, MI-2 (20mg/kg), sorafenib (30mg/kg) or combination treatment for 14 days. ***P* < 0.01, ****P* < 0.001. **i,** Representative images of H&E, GPX4 and 4-HNE staining performed on formalin-fixed paraffin-embedded OSRC-2 xenografts from indicated groups. **j,** Tumor volumes of LoVo xenografts in BALB/c nude mice were measured following vehicle, MI-2 (20 mg/kg), regorafenib (5mg/kg) or combination treatment for 14 days. **P* < 0.05, ****P* < 0.001. **k,** Representative images of H&E, GPX4 and 4-HNE staining were performed on formalin-fixed, paraffin-embedded LoVo xenografts from indicated groups.
