## Supplementary figures and images for "Identification of a Novel Ferroptosis Inducer with Dual Modulatory Effects on GPX4 Activity and Stability"

### Extended Data Fig 1

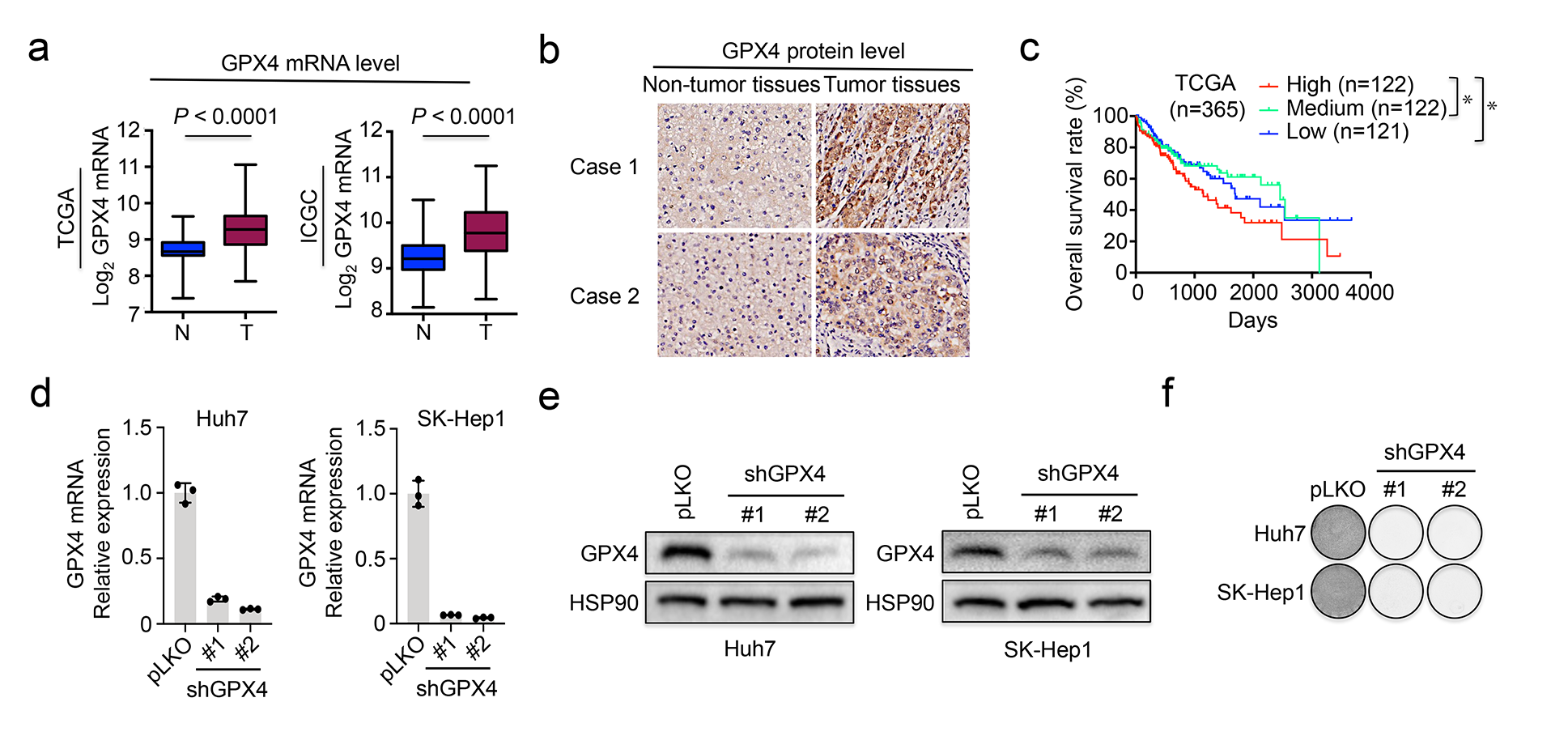

### Extended Data Fig 2

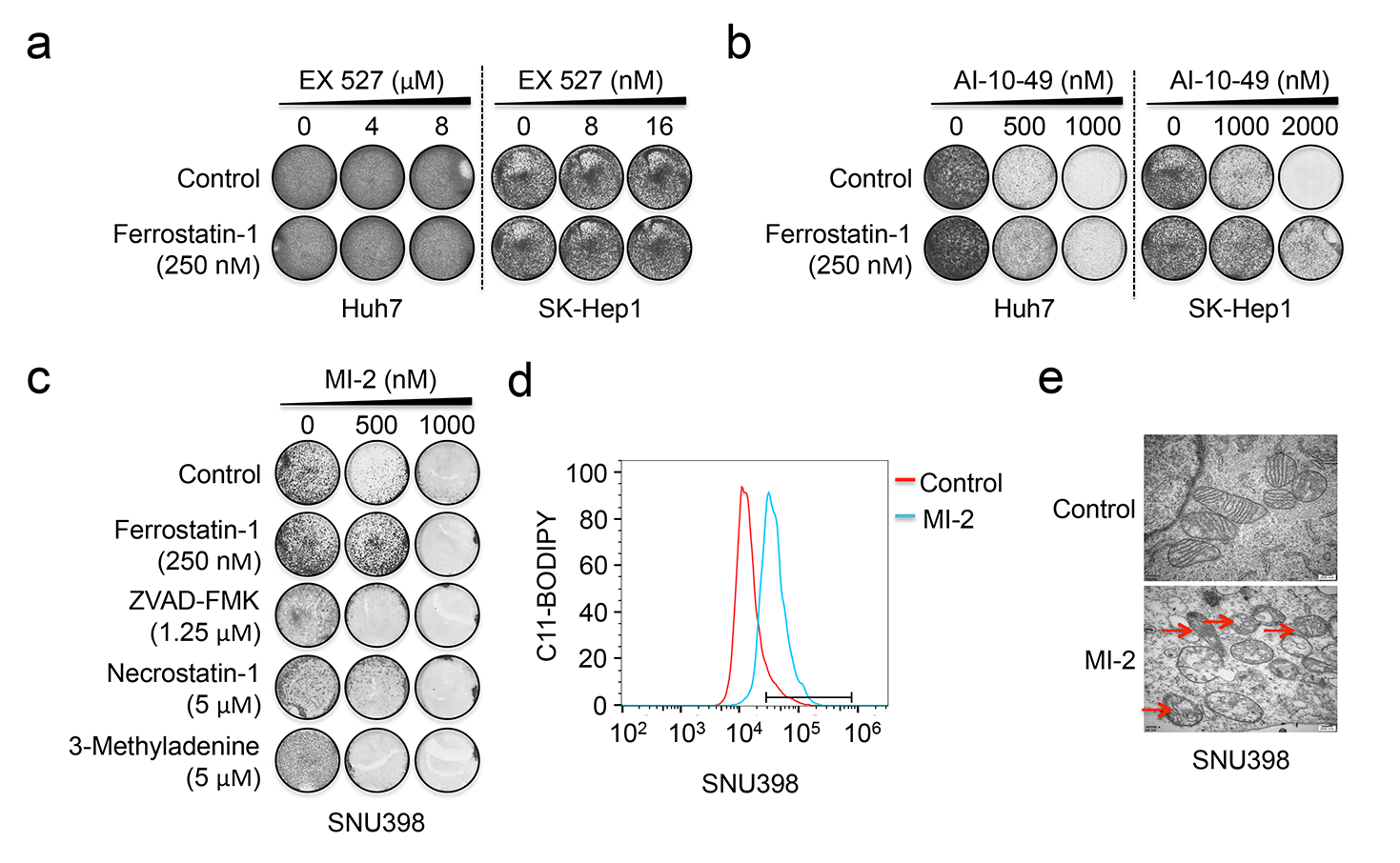

### Extended Data Fig 3

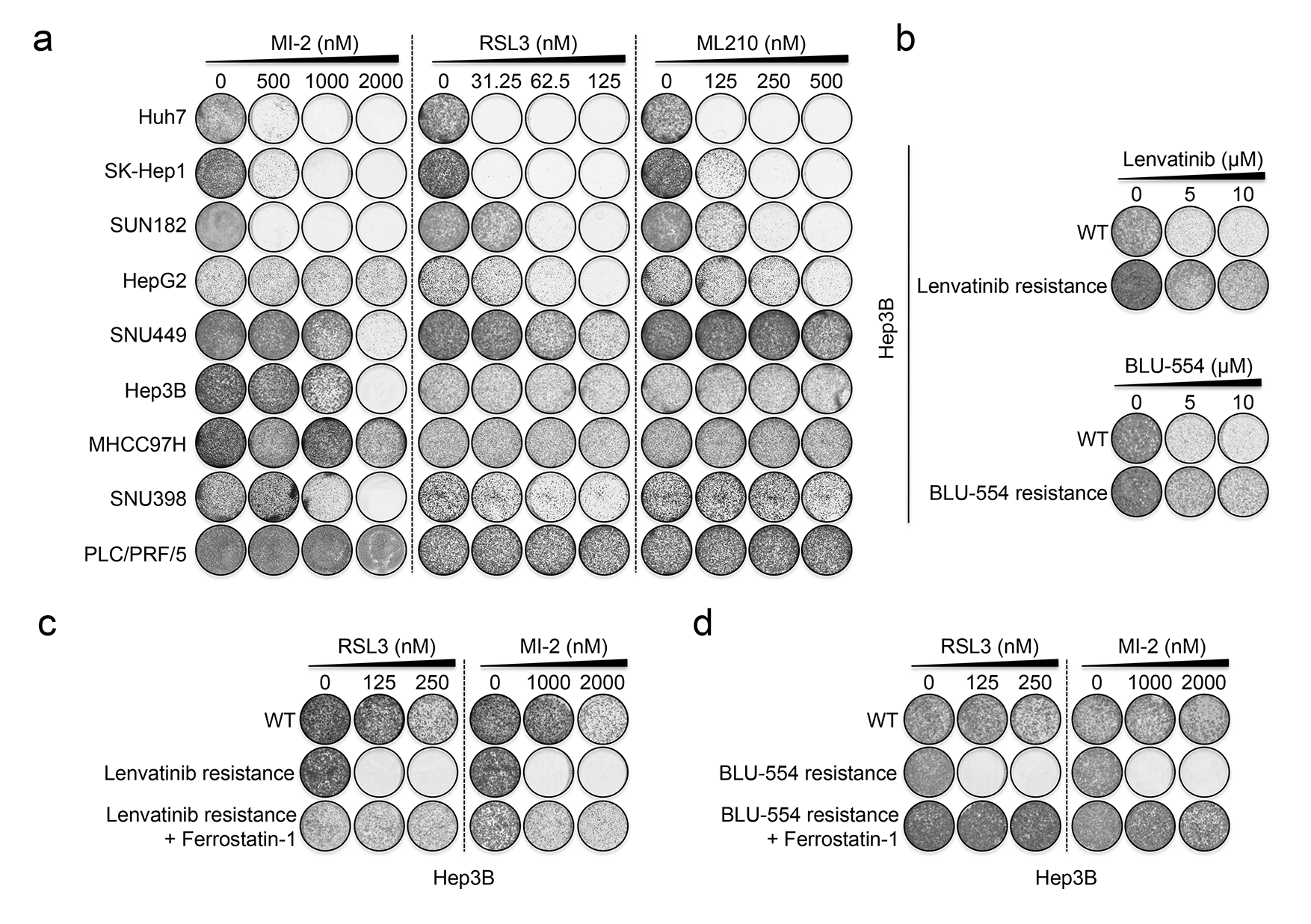

### Extended Data Fig 4

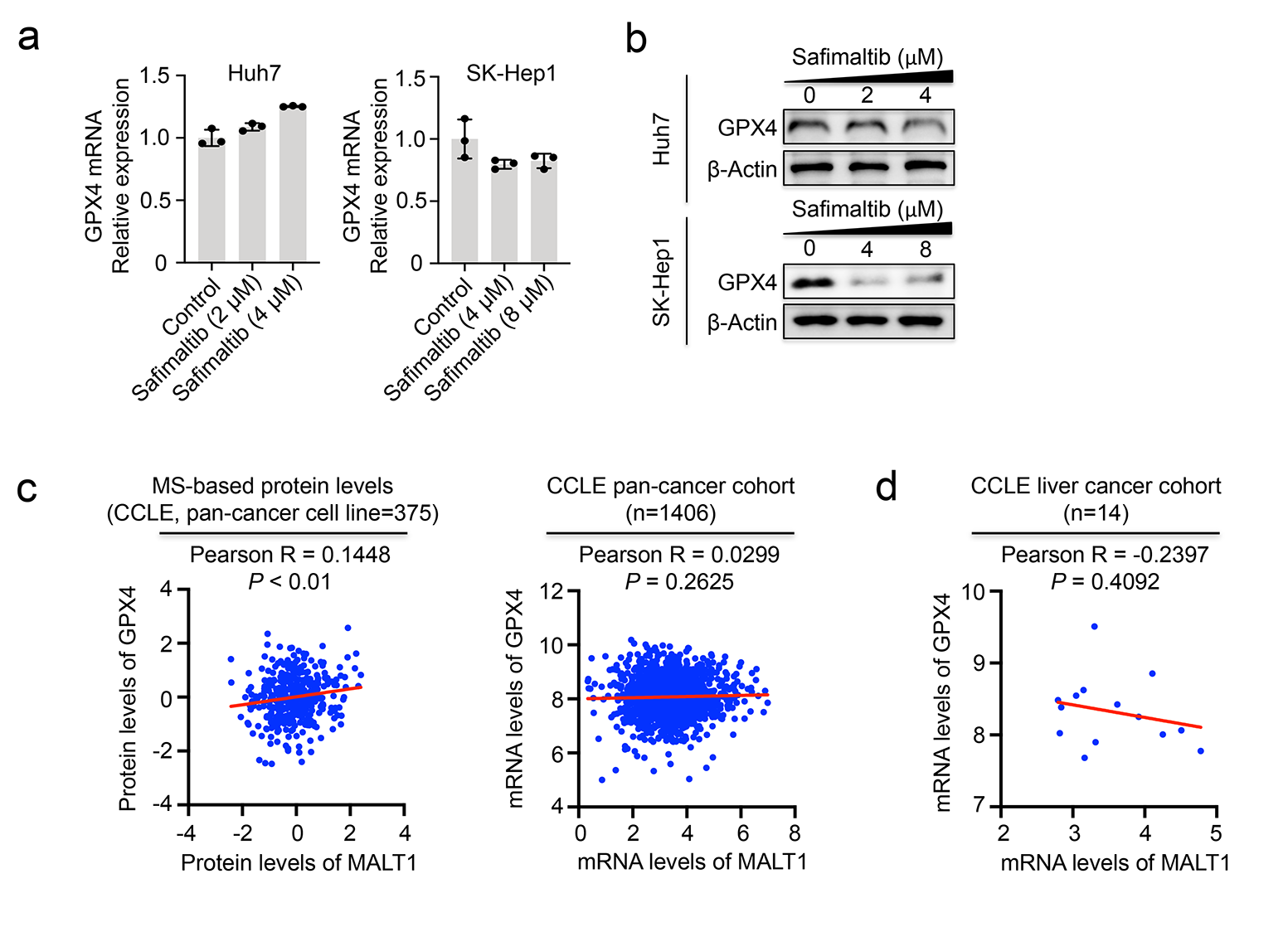

### Extended Data Fig 5

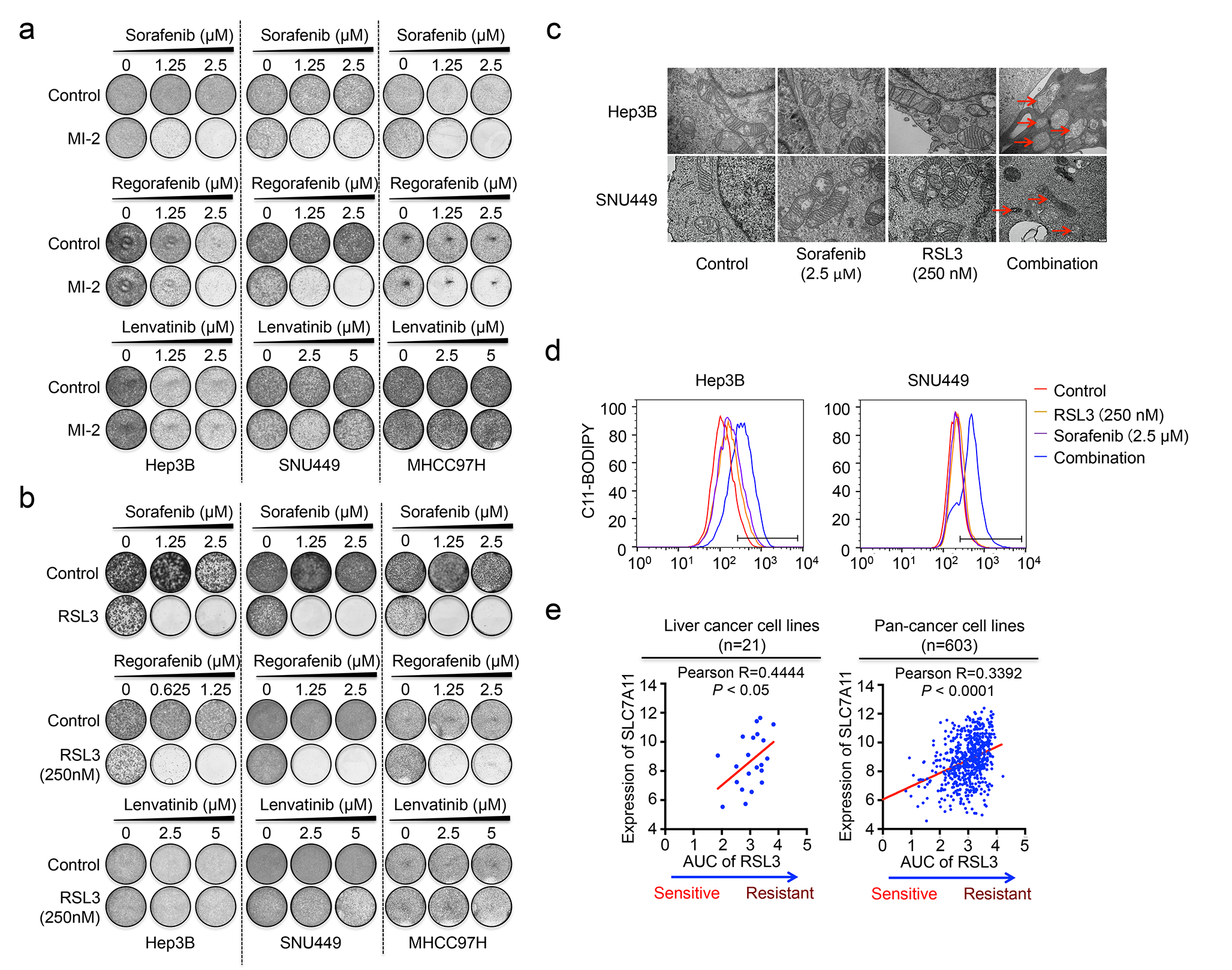

### Extended Data Fig 6

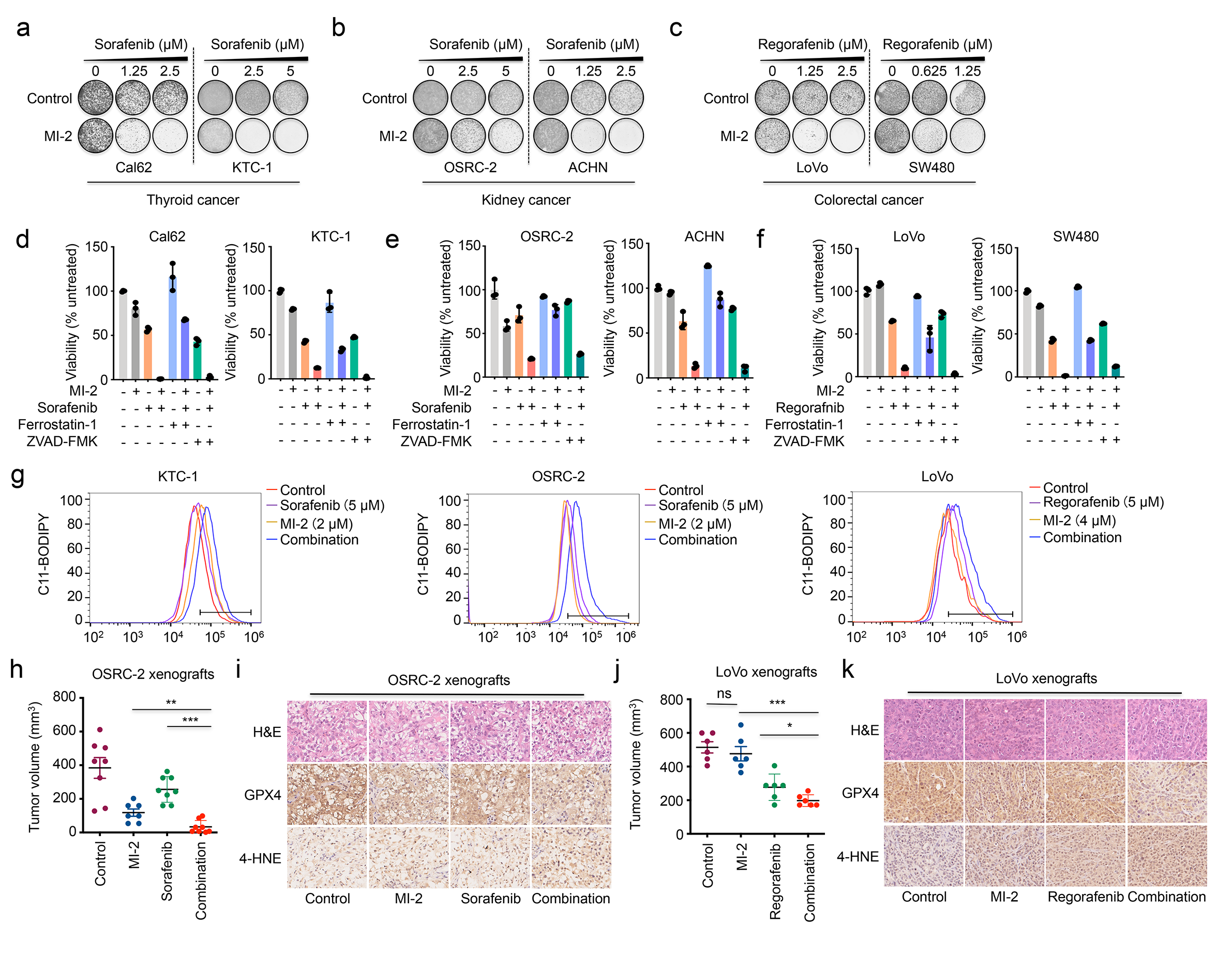
